# HDAC1 regulates the chromatin landscape to establish transcriptional dependencies in chronic lymphocytic leukemia

**DOI:** 10.1101/2020.08.03.232561

**Authors:** Tzung-Huei Lai, Hatice Gulcin Ozer, Pierluigi Gasparini, Giovanni Nigita, Rosario Destefano, Lianbo Yu, Janani Ravikrishnan, Tzung-Lin Tsai, Rosa Lapalombella, Jennifer Woyach, Vinay Puduvalli, James Blachly, John C Byrd, Deepa Sampath

**Affiliations:** Department of Internal Medicine, Division of Hematology, Comprehensive Cancer Center, The Ohio State University, Columbus, Ohio, USA; Department of Biomedical Informatics, The Ohio State University, Columbus, Ohio; Human Cancer Genetics Program, Comprehensive Cancer Center, The Ohio State University, Columbus, Ohio, USA; Center for Biostatistics, The Ohio State University, Columbus, Ohio, USA

## Abstract

HDAC1 is a key regulator of gene expression in cancer. We identified a critical role for HDAC1 in establishing the transcriptional dependencies essential for survival in chronic lymphocytic leukemia (CLL) by profiling HDAC1 with BRD4, H3K27Ac superenhancers, H4K9Ac, chromatin accessibility signatures, Pol2 measurements and expression signatures to generate a regulatory chromatin landscape. Superenhancers marked by high levels of acetylation and BRD4 paradoxically also recruited the highest levels of HDAC1. HDAC inhibition poisoned transcription at these loci to selectively disrupt B-cell transcription factors and B-cell receptor signaling. HDAC1 was also recruited genome-wide at promoters without superenhancers to repress expression; HDAC inhibition induces these genes which include key microRNA networks that reciprocally downregulate CLL specific survival and driver genes. Our work provides a compelling rationale for profiling HDAC1 across cancers to characterize its role in driving the transcriptional dysregulation that is a hallmark of most cancers and develop epigenetic therapeutic strategies.

**Significance:** Our work definitively establishes the composition of the regulatory chromatin that enables HDAC1 to function as an activator and repressor at distinct target genes within the same tumor to drive transcriptional dysregulation and allow the expression of B cell specific signaling and survival networks that are critical for survival.

## Introduction

Chronic lymphocytic leukemia (CLL) is a quiescent disease that exhibits substantial cytogenetic, molecular and clinical heterogeneity^1^. Patients with adverse prognostic markers such as deletions in chromosome 11 or 17, mutations in tumor suppressor, splicing factors or signaling molecules such as TP53, SF3B or NOTCH experience clonal evolution, progressive disease and inferior survival^1^. Despite vast heterogeneity, CLL patients share a common deregulated transcriptional signature that includes the abundant expression and activation of several components of the BCR signaling pathway including BTK^2,3^, anti-apoptotic survival proteins such as Bcl2^4^, loss of expression of key microRNA such as miR-15a and miR-16^5,6^ and a profound immunosuppression with multiple T cell defects^7^. These pathways are critical to the biology of CLL. For instance, loss of the microRNAs miR-15a and miR-16 due to deletions or epigenetic inactivation are linked to the genesis of CLL in humans and mouse models and high levels of the anti-apoptotic survival proteins Bcl2 and Mcl1 ^5,6,8^. Inhibiting BTK with small molecule inhibitors of such ibrutinib or acalabrutinib elicited sustained and durable clinical responses all CLL patients across prognostic subgroups to become a standard of care in the US^9–11^. However, complete responses are rare and therapy-associated mutations lead to therapy resistance with poor outcomes. Alternative therapies such as venetoclax that target Bcl2 offer clinical benefit to patients with ibrutinib resistant disease^12,13^. Similar to BTK inhibitors, mutations in Bcl2 drive progression on venetoclax^14^ with poor survival outcomes. Similarly, therapy with genetically engineered T cells (CART) to circumvent T cell defects in CLL have elicited complete responses but have limited longevity^15,16^; Since CLL survival depends on transcriptionally driven oncogenic networks^17^, we undertook a comprehensive evaluation of the regulatory chromatin landscape that controls transcriptional dependencies in order to identify rational new targets.

Transcription is largely controlled by acetylation writers such the histone acetyltransferases (HATs), erasers such as the histone deacetylases (HDACs) and readers such as BRD4 which dictate chromatin accessibility to transcription factors and RNA Pol2 to regulate gene expression^18^. The eraser, histone deacetylase HDAC1 has been widely studied as a transcriptional repressor due to its ability to deacetylate histones, compact chromatin and prevent transcription^19^. High HDAC enzymatic activity was associated with poor overall survival in CLL^20^. As a repressor, HDAC1 helps to establish essential dependencies in CLL by driving the epigenetic silencing of several microRNAs (miRs) such as miR-210, miR15-a, miR16 and miR-29 to lead in part to elevated expression of BTK^2^ as well as the key survival proteins Bcl2 and Mcl1^8^. miRs constitute an important regulatory layer that control the transcriptome and disease state by post-transcriptional mechanisms^21^.

Superhancers are large regulatory elements characteristic of cancer cells that are densely occupied by transcription factors, histone marks such as acetylated lysine 27 (H3K27Ac) and the bromodomain-containing protein BRD4^22^. BRD4 drives transcription by recognizing H3K27Ac marks at promoters and enhancers to recruit initiation/elongation complexes and RNA Pol2^23^. In CLL, BRD4 associates with a small set of oncogenic SEs to drive the deregulated expression of genes implicated in CLL pathogenesis and survival^17,22^. Targeting BRD4 causes SEs to collapse leading to a decrease in oncogenic transcription and potent anti-tumor activity and highlights the dependence of cancer cells on SE driven transcription^17,22^. Paradoxically, in rhabdomyosarcoma cell lines, levels of HDAC1 were bound at SEs^18^ and facilitated transcription by driving elongation^24,25^. In parallel, HDACs were shown to regulate BRD4 availability in cell lines as HDAC inhibition trapped BRD4 at hyperacetylated sites, decreasing its availability to induce transcription at non-hyperacetylated promoters.^25^

Together, these reports identify numerous unanswered questions of relevance in CLL. Since BRD4 was already shown to associate with SEs in CLL it is unknown whether HDAC1 becomes recruited at the same chromatin loci as BRD4 at SEs and TEs in CLL. Studies on whether HDAC1 functions as a transcriptional activator or repressor or both are lacking. Whether HDAC inhibition is sufficient to disrupt BRD4 driven oncogenic transcription is not known. Additionally, although CLL is characterized by widespread dysregulation of miRs which impact survival and progression^2,5,8,26^, the role of HDACs in miR dysregulation in CLL is poorly known. Finally, the importance of HDAC1 in the regulatory landscape of CLL and in establishing the transcriptional dependencies in CLL is unknown.

Here, we show that the composition of the HDAC1-containing regulatory chromatin controls transcriptional dysregulation in CLL. We profiled HDAC1, BRD4, enhancer and promoter specific histone modifications and RNA Pol2 and functionally integrated these with chromatin accessibility and target expression signatures to identify three dominant regulatory patterns in the CLL genome based on the composition of the HDAC1-containing regulatory chromatin. Cluster I consisted of high levels of HDAC1 extensively co-recruited with BRD4 across SEs at a specific group of protein coding and microRNA genes. Despite high levels of HDAC1 these sites also displayed high levels of acetylation at its histone target, H3K9Ac, open chromatin signatures, robust RNA Pol2 engagement and high transcript expression indicating that HDAC1 in this configuration of the regulatory chromatin predominantly facilitated gene expression. Rank order of the genes at HDAC1 associated BRD4 dense SE identified a number of genes important to survival, BCR signaling and immune dysfunction in CLL. HDAC inhibition within this subset significantly reduced SE and H3K9Ac signals to cause spreading of BRD4 enhancers, and reducing RNA Pol2 initiation to selectively halt the expression of B cell specific transcription factors, molecules implicated in immune dysfunction, survival and signaling proteins. Cluster II, consisted of focal recruitment of HDAC1 to sites not marked by SEs or BRD4 but still having high levels of H3K9Ac, open chromatin signatures and Pol2 recruitment at sites. The bulk of transcripts in this subset had low to moderate expression indicating that HDAC1 functioned to limit the expression of target genes within this cluster. A few notable genes that became upregulated after HDAC inhibition included miRs that targeted BTK and compromise survival in CLL^27^. Cluster III harbored promoters that recruited HDAC1 alone. These were unmarked by SEs, BRD4, had low or no H3K9Ac, lacked open chromatin signatures and RNA Pol2 and represented gene clusters that underwent HDAC1-mediated transcriptional repression. This subset included multiple microRNA genes regulated critical B cell drivers such as IKZF3, BTK, and SYK. Thus, systematic profiling of the HDAC1-containing regulatory chromatin establishes the mechanism by which HDAC1 functions as a facilitator and repressor of gene expression at distinct targets within tumors to drive transcriptional dysregulation and facilitate survival. HDAC1 profiling also uncovers a transcriptional vulnerability since HDAC inhibition not only epigenetically abrogated BRD4 driven BCR and survival signaling it also activated microRNA networks that converged on targeting driver genes in CLL.

## Results

### The composition of the HDAC1 containing regulatory chromatin establishes the transcriptional state of the CLL genome

This study used a cohort of primary CLL tumor cells from 57 patients that represented clinically relevant prognostic subtypes including immunoglobulin-heavy chain (IgVH) mutational status, FISH, cytogenetics, and presence or absence of prior therapy as described in **Table S1**. We have previously shown the over expression of BRD4 in CLL samples^17^, we now show using healthy B cells (n=8) and CLL cells (n=16) that both HDAC1 and BRD4 are overexpressed in CLL (**Figure 1A**).

**Figure 1.**
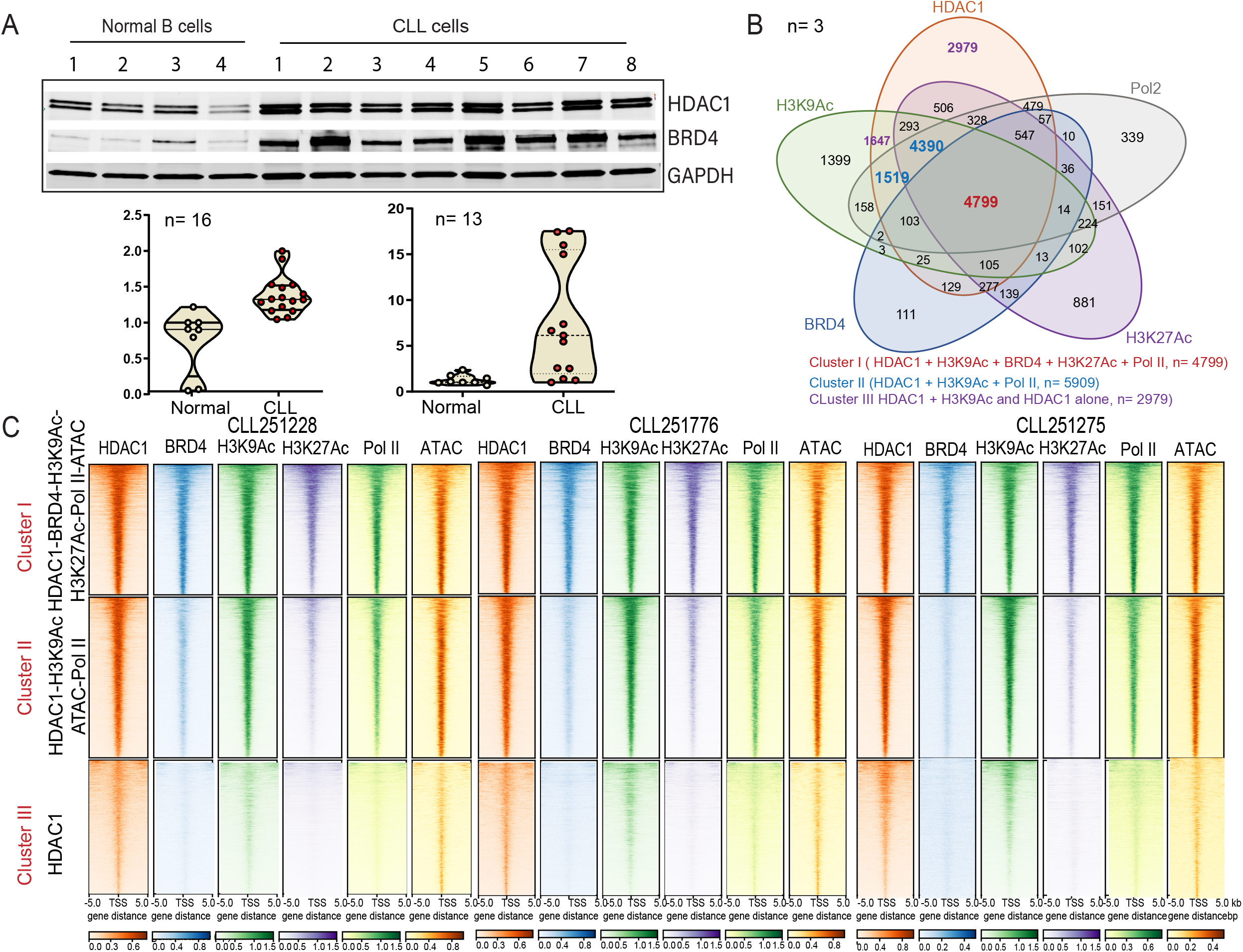
The functional chromatin landscape of CLL as defined by HDAC1 recruitment. **(A)** Expression of HDAC1 and BRD4 in healthy B cells and patient CLL cells are shown in the upper panel and expression levels for HDAC1 and BRD4 from healthy B cells (n=8) and CLL cells (n=16) were quantitated in the lower panels. **(B)** Venn diagram showing the intersection of HDAC1 with BRD4, H3K9Ac, H3K27Ac, ATAC Seq, RNA Pol II recruitment in 3 primary CLL samples; Cluster I (HDAC-BRD4-H3K9Ac-H3K27Ac-ATAC-RNA Pol II – 4799 peaks representing 30% of the transcriptome), Cluster II (HDAC1-H3K9Ac-ATAC-Pol II – 5909 peaks representing 33% of the transcriptome), Cluster III (HDAC1 only – 2979 peaks representing 6 % of the transcriptome). **(C)** ChIP-Seq densities ranked in decreasing order for the genome wide combined occupancy signals of HDAC1 with BRD4, H3K9Ac, H3K27Ac, open ATAC signatures and RNA Pol II engagement (Cluster I), HDAC1 co-recruited with H3K9Ac, ATAC and RNA Pol II (Cluster II) and HDAC alone (Cluster III) centered ± 5 kb of the transcriptional start site (TSS) window in three CLL samples (CLL251288, CLL251766 and CLL251275).

Next, we postulated that the composition of the HDAC1-containing regulatory chromatin controlled the transcriptional dysregulation in CLL. To evaluate this, we conducted chromatin immunoprecipitation (ChIP) linked to deep sequencing to generate the recruitment profiles of HDAC1 across the CLL genome and then integrated it with the binding profiles of BRD4, H3K27Ac SEs and H3K9Ac along with functional measures of transcription such as chromatin accessibility signatures, RNA Pol2 recruitment and gene expression profiles across all HDAC1 bound gene targets. Based on the observed recruitment signatures for the above proteins, we defined three dominant regulatory patters as shown in three CLL samples (CLL251275, CLL251776, CLL251228). Cluster I consisted of extensive and high levels of HDAC1 at chromatin loci that also harbored BRD4, H3K9Ac, H3K27Ac and RNA Pol2 at 4799 peaks close to the transcriptional start sites (TSS) of 5122 genes and represented 30% of the CLL transcriptome (**Figure 1B, 1C and Table S2**). These sites were characterized by hyper-accessible chromatin signatures as determined by ATAC Seq and robust RNA Pol2 recruitment. Cluster II consisted of HDAC1 bound in association with H3K9Ac and RNA Pol2 without BRD4 or SEs at 5909 peaks near the TSS of 5676 genes and accounts for 33% of the transcriptome. These loci also demonstrated hyper accessible chromatin signatures but far less RNA Pol2 recruitment (**Figure 1B, 1C and Table S2**). Cluster III had HDAC1 bound without BRD4, H3K9Ac (low or none) or H3K27Ac at 4626 peaks near the TSS of 1077 genes and accounts for 6% of the transcriptome. This group did not have discernable open chromatin signatures or RNA Pol2 associated at target promoters. (**Figure 1B, 1C and Table S2**). Thus, the presence of HDAC1 in combination with specific chromatin modifiers and modifications establishes the degree of transcriptional permissivity of the CLL genome.

### Dynamics of transcription based on whether HDAC1 associates with SEs and TEs in CLL

Recent reports in cell lines suggest that HDAC1, when recruited to core or super enhancers, positively regulates transcription. Our study identified H3K27Ac peaks at 11,745 (CLL251228), 13,762 (CLL251766) and 14,293 (CLL251275) loci genomewide of which BRD overlapped H3K27Ac at 9213 (CLL251228), 7973 (CLL251776) 4814 (CLL251275) peaks indicating a 95%, 96 and 91% overlap between BRD4 and H3K27Ac (**Suppl Fig 1**). In agreement with previous reports, we identified dense BRD4 loads at a small number of SEs close to 382 (CLL251228), 312 (CLL2501776) and 245 (CLL250275) genes with critical roles in CLL^17,22^. Significantly, we observed extensive and dense loads of HDAC1 at the majority of the BRD4 dense SEs (335/382 (CLL251228), 284/312 (CLL2501776) and 215/245 (CLL250275)) representing an 88%, 91% and 88% overlap between HDAC1 and BRD4 at SEs (**Figure 2A-in red**, **Table S3**). Rank ordering of HDAC1 enriched, BRD4 dense enhancers (**Table S4**) identified multiple genes critical to the pathology of CLL such as those that favored immune dysfunction CXCR4, IL16, and IL4, survival proteins such as Bcl2, core transcription factors in CLL such as Pax5, IKZF1 and IKZF3 and B cell signaling pathway kinases such as BLK among the top enhancers (**Figure 2B**).

**Figure 2.**
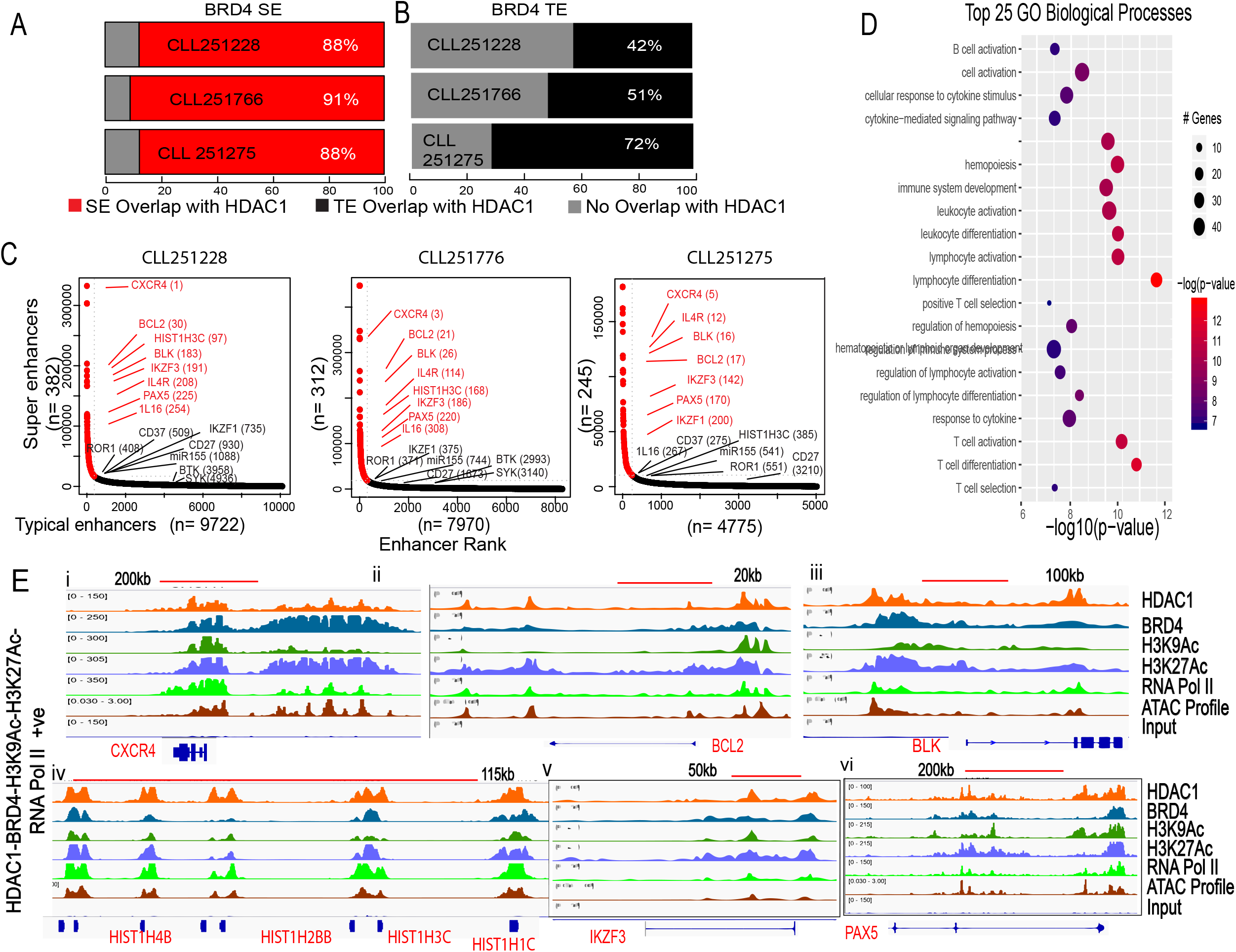
Distinct HDAC1 linked SE and non-SE profiles in CLL. **(A)** Of 382 (CLL251228), 312 (CLL2501776) and 245 (CLL250275) genes with BRD4 dense marks at SEs, 335 (88%) (CLL251228), 284 (91%)(CLL2501776) and 215 (88%)(CLL250275) also strongly bound HDAC1. **(B)** Of 4775 (CLL251228) 7970 (CLL251776), and 9772 (CLL251275) genes that had BRD4 associated with typical enhancers (TEs) 3423 (71% - CLL250228), 4096 (51% - CLL250776) and 4110 (42% - CLL 250275) also co-recruited HDAC1 **(C)** Recruitment of HDAC1 at enhancer profiles of three representative CLL samples. The intersection of CHIP-Seq signals of HDAC1, BRD4 and H3K9Ac SEs were generated after which Enhancers were then rank ordered by their BRD4 load in association with HDAC1 and harbored 382, 312, 245 BRD4 peaks at SEs and 8441, 14468 and 17201 BRD4 peaks at typical enhancers (TEs) in CLL251228, CLL2501776 and CLL250275. Selected genes bound by HDACs at BRD4 linked SEs among the top enhancers are marked in red and selected genes bound by HDACs with BRD4 at TEs are marked in blue. SE and TE identification for all samples are listed in Table S4. **(D)** TOPP Gene Suite was used to derive functional annotations of the top biological processes associated with HDAC bound at BRD-SEs. Top 25 pathways (based on z-scores|) associated with the genes are shown and color coded based on specificity from orange to blue and while the size reflects the number of genes that make up the category. **(E)** Overlay of HDAC1, BRD4, H3K9Ac, H3K27Ac, ATAC-Seq profiles and RNA Pol II densities at selected top genes showing HDAC association at BRD4 SEs in Cluster I (i-vi - CXCR4, Bcl2, BLK, HIST1H3A/B/C, Pax5, IKZF3. Each track is representative of the ChIP-Seq binding densities from three independent CLL patient samples.

In addition to its association at SEs, HDAC1 was also recruited with BRD4 at 3423 of 4775 (71%, CLL251228) 4096 of 7970 (51%, CLL251776), and 4110 of 9772 (42% CLL251275) at core enhancers (TEs) (**Figure 2A–in black,Table S3**). We investigated several of these genes if they represented the top genes associated with TEs or represented components of the signaling pathways that make up the top enhancers (SE) in CLL. Based on these criteria and we identified the B cell transcription factors such as ROR1, and IKZF1, B cell signaling kinases such as BTK and SYK, and miR155, CD37 and CD27 among the top relevant genes that bound HDAC1 at TEs (**Figure 2B**). Comprehensive functional annotation of the top genes in Cluster I (that represented HDAC1dense loci bound at BRD4 containing TEs and SEs) identified B cell activation, lymphocyte activation and several immune pathways among that the top biological processes (**Figure 2C**). The binding profiles of select genes from Cluster I illustrating high levels of HDAC1 load associated with BRD4 SEs (**Figure 2D, i-vi, Suppl. Figure 2A,i-ii**) and TEs (**Suppl. Figure 2Aiii-ix**) are shown. Expression analysis of select genes bound by HDAC1 at BRD4 SEs and TEs (Cluster I) demonstrated that these genes were expressed at levels equal to or higher than that found in healthy B cells both at the transcript (**Figure 3A, i-viii and Supp. Figure 2B i-ii**) and protein levels (**Figure 3B**). These findings indicate that HDAC1 when co-recruited with BRD4 at SEs and TEs positively facilitate the transcription of target genes, many of which are critical in the pathobiology of CLL.

**Figure 3.**
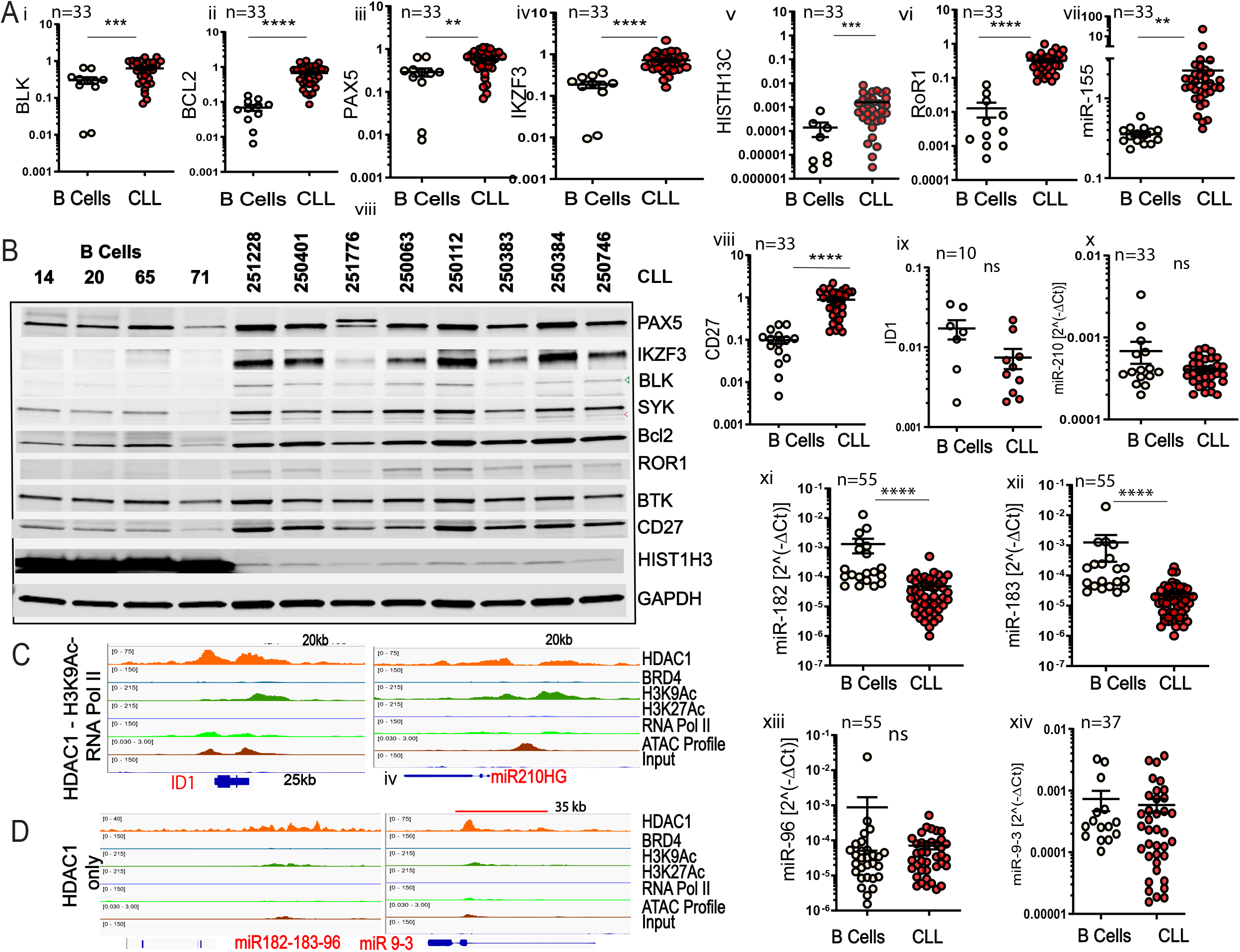
HDAC1 associated at SEs and TEs facilitates target gene expression. **(A)** mRNA expression levels of the genes whose tracks were shown in Figure 2E and Supp. Figure 3; Cluster I ([i-viii]-Blk, Bcl2, Pax5, IKZF3, HIST1H3C, ROR1,miR-155, CD27), Cluster II ([ix-x]-ID1, miR210) and Cluster III [(xi-xiv]-miR-182 and miR9-3) in 33 CLL samples compared to 11-15 heathy B cell samples except ID1 (CLL=10, B cells=7). *,**,***,**** p<.5,.01,.001,.0001, Student T-test, Graph pad software. **(B)** Protein expression levels of genes evaluated in (A) in eight CLL samples compared to healthy B cells. **(C**) Overlay of HDAC1, BRD4, H3K9Ac, H3K27Ac, ATAC-Seq profiles and RNA Pol II densities at selected genes showing HDAC association with H3K9Ac, open ATAC signatures and RNA (Cluster II, ID1, mir-210) **(D)** Overlay of HDAC1, BRD4, H3K9Ac, H3K27Ac, ATAC-Seq profiles and RNA Pol II densities at selected genes which recruit HDAC1 only (Cluster III) (miR-182, miR9-3).

We then evaluated genes that displayed high levels of H3K9Ac and Pol2 recruitment and had open chromatin signatures despite dense HDAC1 recruitment at their promoters (Cluster II). The binding profiles of 2 representative genes ID1 and miR210 are shown (**Figure 3C**). Both ID1, and miR-210 showed lower expression levels compared to healthy B cells (**Figure 3A ix-x**) and ID1 protein was undetectable (not shown). These results indicate that suggesting that when the regulatory chromatin comprised of HDAC1, despite high levels of its deacetylation target H3K9Ac and Pol2 engagement at the same targets, HDAC1 recruitment served to limit the expression of genes in Cluster II. Finally, genes that recruited HDAC1 with low or no H3K9Ac and without any other regulatory modifications (Cluster III) had closed chromatin signatures, no RNA Pol2 recruitment (Figure 3D) and largely (>80%) had low to moderate expression. These genes encompassed several key microRNA genes (**Figure 3A, xi-xiv**). Under, this chromatin configuration, HDAC1 functioned as a transcriptional repressor.

### HDAC inhibition causes enhancer spreading and local loss of BRD4 at HDAC1 bound SEs and TEs to selectively modulate CLL driver genes

High levels of HDAC at key loci across the genome may uncover a selective transcriptional vulnerability. Therefore, we exposed CLL cells to abexinostat, an HDAC inhibitor that targets HDAC1 with greater potency than other HDACs^28^, for 6 h to inhibit HDAC function, before conducting ChIP to identify changes in the binding profiles of HDAC1, H3K9Ac, BRD4, dynamics of SEs and TEs, and RNA Pol2. Changes in recruitment patterns of the regulatory chromatin were integrated with changes in the expression levels of genes located within the boundaries of the HDAC1 containing chromatin across all clusters (Cluster I-III) followed by a detailed evaluation of each cluster.

We began our analysis with genes that strongly recruited HDAC1 at BRD4 dense SEs (Cluster I) In rhabdomyoma cell lines, most of the HDACs were associated with SEs and HDAC inhibition caused a collapse of SEs, BRD4 spreading, loss of RNA Pol2 and a loss of SE driven transcription^18,25^. Similarly loss of transcription after BRD4 inhibition depended on disrupting SE architecture^17,22^. However, our study in primary tumor cells showed that HDAC inhibition caused a significant decrease in HDAC1 binding and contrarily, decreases in H3K9Ac and H3K27Ac binding (**Figure 4A**) suggesting the establishment of a non-permissive chromatin at these target genes. This was accompanied by a disruption of BRD4 peaks which underwent diffusion and spreading (**Figure 4A-4D**) and likely reflects a loss of functional enhancer activity. However, when we related decreases in BRD4 and H3K9Ac binding to chromatin accessibility and RNA Pol2 recruitment we found a bimodal signal where the decrease in ATAC accessibility signature and RNA Pol2 recruitment (**Figure 4A**) at a large set of driver genes was counteracted by increase in chromatin accessibility and RNA Pol2 from a small set of genes that became induced leading to an apparently stable ATAC and RNA Pol2 profile without net losses or gains. These changes are quantitated in **Table S5**.

**Figure 4.**
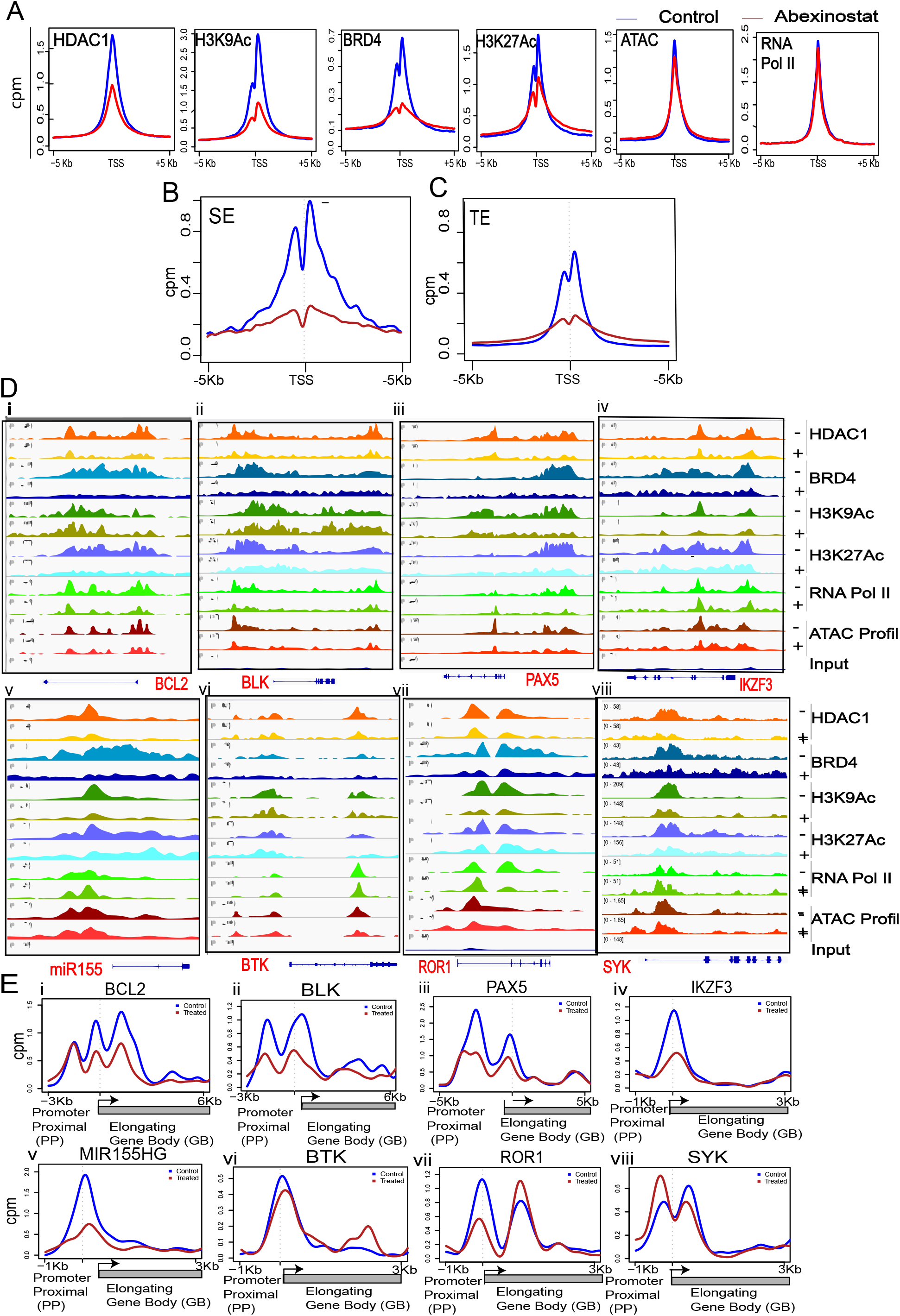
HDAC inhibition disrupts BRD4 and RNA Pol II at SEs. **(A)** Changes in the HDAC1, BRD4, H3K9Ac, H3K27Ac ATAC Seq and RNA Pol II ChIP-Seq meta-gene profiles before (blue) and after exposure (red) to the HDAC inhibitor, abexinostat for 6h at all 4799 peaks representing the union of HDAC1 co-recruited with BRD4 at H3K27Ac enhancers (SEs and TEs), H3K9Ac (transcriptionally permissive chromatin) and Pol II with open ATAC signatures (Cluster I). Normalized tag counts for the HDAC1, BRD4, H3K9Ac, H3K27Ac and Pol II before and after HDAC inhibition are supplied in Supplementary Table S5 **(B and C)** Effect of HDAC inhibition on BRD4 peak counts at SEs and TEs. BRD4 ChIP-Seq meta gene profiles showing the change in BRD4 enrichment across H3K27Ac SEs peaks and at TEs before (blue) and after exposure (red) to the HDAC inhibitor, abexinostat for 6h at all genes that had HDAC1 co-recruited with BRD4 at SEs and TEs (Cluster I). **(D)** Genome browser tracks showing the change in HDAC1 (orange), H3K9Ac (green), BRD4 (teal),H3K27Ac (dark Blue), RNA Pol II (green), ATAC Seq peaks (burgundy) in a representative CLL sample out of three at baseline and after exposure to abexinostat for 6 h, HDAC1 (yellow), H3K9Ac (navy), BRD4 (olive),H3K27Ac (sky Blue), RNA Pol II (dark green), ATAC Seq peaks (red) at each of the genes that bound HDAC1 at BRD4 dense SEs (Bcl2, BLK, Pax5, IKZF3) and TEs (miR-155, BTK, Syk and Ror1). **(E)** Effect of HDAC inhibition on RNA Pol II initiation at promoters. RNA Pol II ChIP-Seq **m**eta gene profiles showing changes in reads per million (rpm) for RNA Pol II occupancy from −5 to - 1KBkb of the proximal promoter (PP) centered around the transcriptional start site (TSS) to +3 to +6 KB of the gene body (GB) at each of the genes shown in (E) in CLL representative samples before (blue) and after exposure to abexinostat for 24 h (red).

To investigate this apparent discord between layers of epigenetic modifications such as the loss of HDAC1, H3K27Ac and BRD4 binding after drug exposure and lack of corresponding losses in open ATAC signatures and RNA Pol2 binding, we focused on genes that lost expression after HDAC inhibition in Cluster I (**Table S6**). We confirmed that HDAC inhibition largely decreases the BRD4 load at SEs (**Figure 4B**) and TEs (**Figure 4C**) and found that CLL cells bound twice as much BRD4 when associated with SEs compared to TEs under baseline conditions. We show the binding patterns of HDAC1, BRD4, H3K27Ac, H3K9Ac, ATAC signatures and RNA Pol2 engagement at selected CLL specific driver genes located close to HDAC bound SEs such as Bcl2, BLK, Pax5, IKZF3 (**Figure 4D,i-iv**) and IL4R, CXCR4, IL16, CD27, CD37 (**Supp. Figure 3**) and TEs such as miR-155, BTK, Ror1 and Syk (**Figure 4D,v-viii**) after HDAC inhibition. The chromatin binding profiles at these CLL specific driver genes demonstrate loss of binding of HDAC1, decrease in H3K27Ac and H3K9Ac signals, BRD4 spreading, decrease in open ATAC signatures and RNA Pol2 engagement. To determine whether the reduced RNA Pol2 engagement was due to decrease in initiation or due to RNA Pol2 pausing we quantitated the amount of Pol2 at the TSS compared to the gene body and determined that was a significant reduction in the occupancy of RNA Pol2 at transcription initiation at the proximal promoter for Bcl2, BLK, Pax5, IKZF3, miR155 and ROR1 but not at BTK or SYK (**Figure 4E, i-viii**).

Inhibitory changes in the regulatory chromatin resulted in steep declines in the transcript levels of a number of CLL specific core transcription factors and driver genes by RNA Seq in 10 CLL samples exposed to abexinostat for 24h. Fold change rank ordering of differentially expressed genes allowed the selection of 422 genes that showed a greater than 4-fold decrease or increase in with a 10% false discovery rate. This set included Pax5, IKZF3, BLK, BTK, Bcl2, miR155, IL16, IL4R, Ror1, Syk, IKZF1, Mcl-1, CD37 and CD27 (**Figure 5A, Table S6**). At baseline over 80% of the genes in Cluster I were expressed at high levels; HDAC inhibition caused a downregulation in 75% of these genes (**Figure 5B**, in orange). Validation of RNA Seq results using expression assays confirmed the declines in BCL2, BLK, Pax5, IKZF3, miR155, Ror1, Syk, CD27, IL4R, CD37, IL16 and CXCR4 expression in all 10 CLL samples exposed to abexinostat for 24 h (**Figure 5C, Supp. Figure 4A**). Loss in transcript levels was mirrored by decreases in in the levels of the cognate proteins; we show a few representative proteins such as Pax5, BTK and IKZF3 that were exquisitely sensitive to HDAC inhibition and showed the steepest declines in protein levels (**Figure 5D and Supp. Figure 4B**).

**Figure 5.**
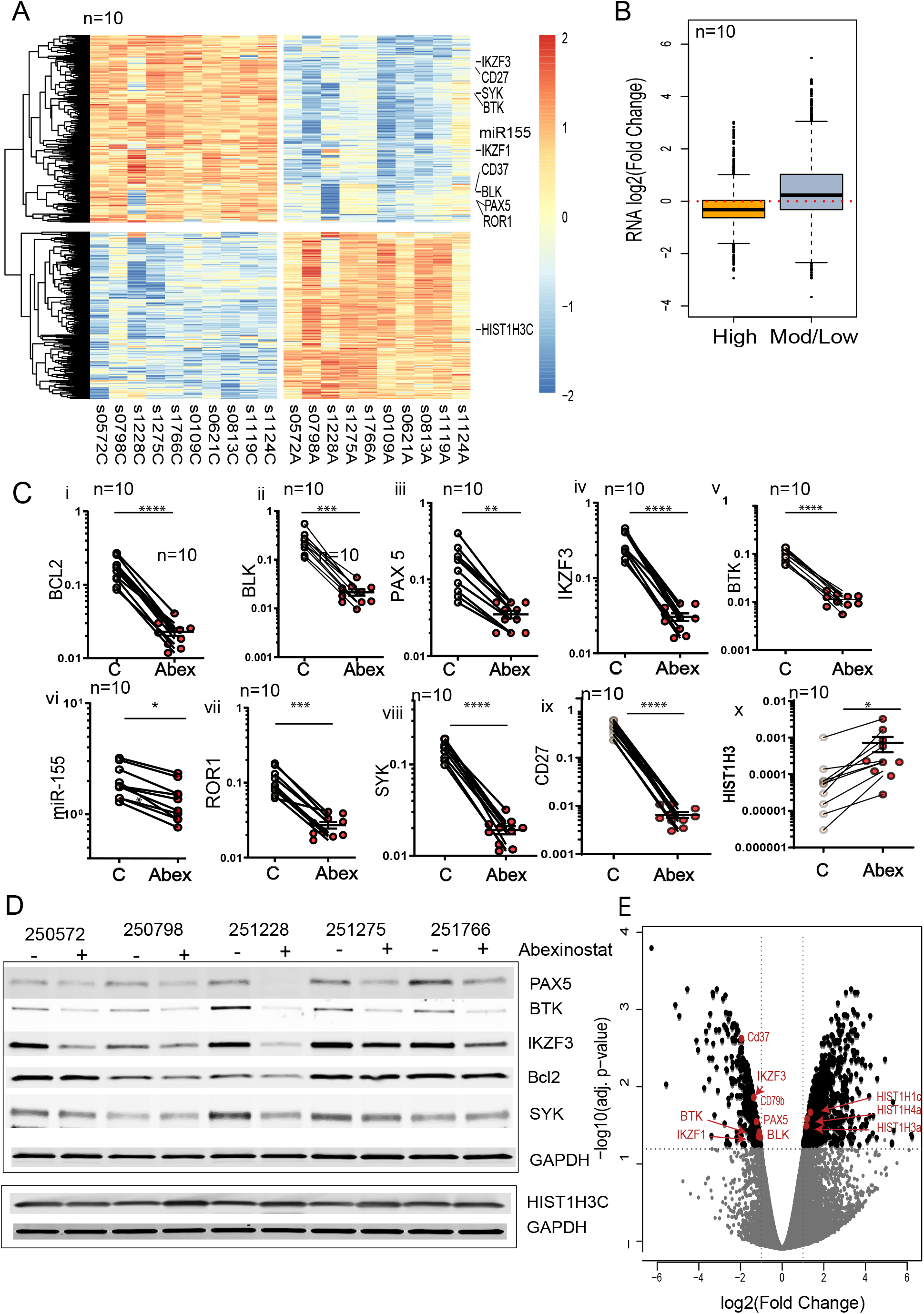
Dynamics of transcription after HDAC inhibition. **(A)** RNA Seq of 10 CLL samples treated with the HDAC inhibitor abexinostat (0.4 μM, 24h). Change in mRNA transcripts was assessed by RNA Seq compared with control samples; top 200 genes that showed a Log 2-fold change with p<0.05 of a total of 1000 protein coding genes that had HDAC bound across SEs and TEs **(B)** Based on average transcript expression at baseline, genes that bound HDAC1 across SEs and TEs were stratified into high (80%) or moderate/low (20%) expressers. mRNA Seq was conducted to assess the change in gene expression after exposure to abexinostat for 24h. 75% of transcripts with high expression at baseline showed a decrease; 25% of low expressers were an upregulated. **(C)** Validation of RNA Seq results using real time-PCR expression assays quantitating changes in expression levels of selected genes from Cluster I ([i-x]-Bcl2, BLK, Pax5, IKZF3, BTK,miR-155, Ror1, Syk, CD27 and HIST1H3C) from 10 CLL samples before and after exposure to abexinostat for 24h. *,**,***,**** p<.5,.01,.001,.0001, Student T-test, Graph pad software. (**D**) Change in protein expression of genes in (C) in 5 CLL samples before and after exposure to abexinostat for 24h **(E)** RNA Seq of from Eμ-Tcl1 mice with advanced CLL treated with abexinostat. Tcl1 mice bearing >10% CD19+CD5+ cells leukemic cells were randomized to vehicle or abexinostat administered thrice weekly for 2 weeks. Spleen CLL cells were isolated and subjected to RNA Seq. The profiles were then compared to those obtained in primary CLL cells exposed to abexinostat to identify shared signatures. Key CLL drivers such as IKZF3, Pax5, Blk and BTK among the top 300 genes that showed a significant downregulation (log-2 fold change with p<0.05) in vivo in response to HDAC inhibitor are shown.

The small fraction of genes (25%) that became induced based on fold change rank with a 10% false discovery rate (FDR) (**Figure 5A and 5B**, in blue) included members of the structural histone protein HIST1, HIST2 and HIST3. Analysis of the binding patterns of the chromatin regulators indicated that while HDAC1, BRD4, H3K9Ac binding decreased at these promoters, they nevertheless acquired a more open chromatin signature and increased RNA Pol2 engagement; these increases balanced the decreases in Pol2 recruitment at genes that were downregulated accounting for the observed lack of change in global RNA Pol2 and ATAC signatures as shown in Figure 4B. The binding tracks for HIST1H3C and increases in RNA Pol2 recruitment at the promoter and TSS as a representative gene are shown (**Supp. Figure 4C and 4D**). Despite the large fold-increase in HIST1H3C transcript, overall the transcript levels remained low (**Figure 5C,x**), HIST1H3 protein was detectable at baseline and HDAC inhibition led to a modest upregulation of protein levels indicating that while HDAC inhibition caused notable changes in the regulatory chromatin at the promoters for the HIST genes it was not translated into major differences at the protein level (**Figure 5D**).

Lastly, we evaluated whether our results in primary tumor cells could be validated *in vivo* using a Tcl1-driven mouse model of CLL. Eμ-Tcl1 mice were bearing >10% CD19+/Cd5+ leukemia cells by flow cytometric analysis were randomized to vehicle or abexinostat administered thrice weekly for 2 weeks. RNA extracted from sleep was then subjected to RNA Seq and demonstrate that similar to human CLL exposure to abexinostat reduced expression of several CLL driver genes such as Pax5, IKZF3, BTK and BLK and show increases in HIST genes. (**Figure 5E**). Taken together, our results show that Cluster I harbored multiple CLL specific driver genes that were expressed at high levels, HDAC inhibition was sufficient to cause decreases in SE amplitude at select genes leading to a diffusion of BRD4, loss of RNA Pol2 and downregulation of oncogenic and survival promoting driver genes.

### HDAC inhibition at genes bound by HDAC without associated SEs causes a redistribution of BRD4 to induce silenced genes

As described in Figure 1, HDAC1 was also recruited genome-wide, co-bound with high levels of acetylated H3K9Ac and RNA Pol2 engagement (Cluster II). HDAC inhibition at these genes (Cluster II) caused a reduction in HDAC1 and H3K9Ac, but induced hyperacetylation of H3K27Ac at these gene clusters in agreement with other reports^18^. There also was an accumulation of BRD4 specifically at target promoters in keeping with the canonical role of BRD4 in phosphorylating RNA Pol2 to initiate transcription. Correspondingly, there was an increase in open signatures and RNA Pol2 engagement as shown (**Figure 6A**) and summarized in **Figure 6B**. At baseline, over 60% of genes in Cluster II were expressed at moderate or low levels. HDAC inhibition caused minimal changes in genes that were well expressed at baseline, but the induced the majority of genes with moderate or low levels at baseline (**Figure 6C, Table S7**) indicating that despite H3K9AC, open ATAC profiles and RNA Pol2 engagement at baseline HDAC likely functioned as a rheostat to limit the expression of these genes. We show the HDAC inhibitor induced changes in the regulatory chromatin of two representative genes ID1 and miR-210 (**Figure 6D**). Both genes respond to HDAC inhibition with with robust increases in RNA Poll II engagement both at the TSS and over the elongating gene bodies within 6 h (Figure 6E) leading to an induction of transcript levels as validated in 10 CLL samples within 24 h (**Figure 6F**). Despite robust induction at the transcript level, ID1 protein was poorly detected by immunoblots indicating that the protein coding genes in Cluster II were less likely to play key roles in the pathobiology of CLL.

**Figure 6.**
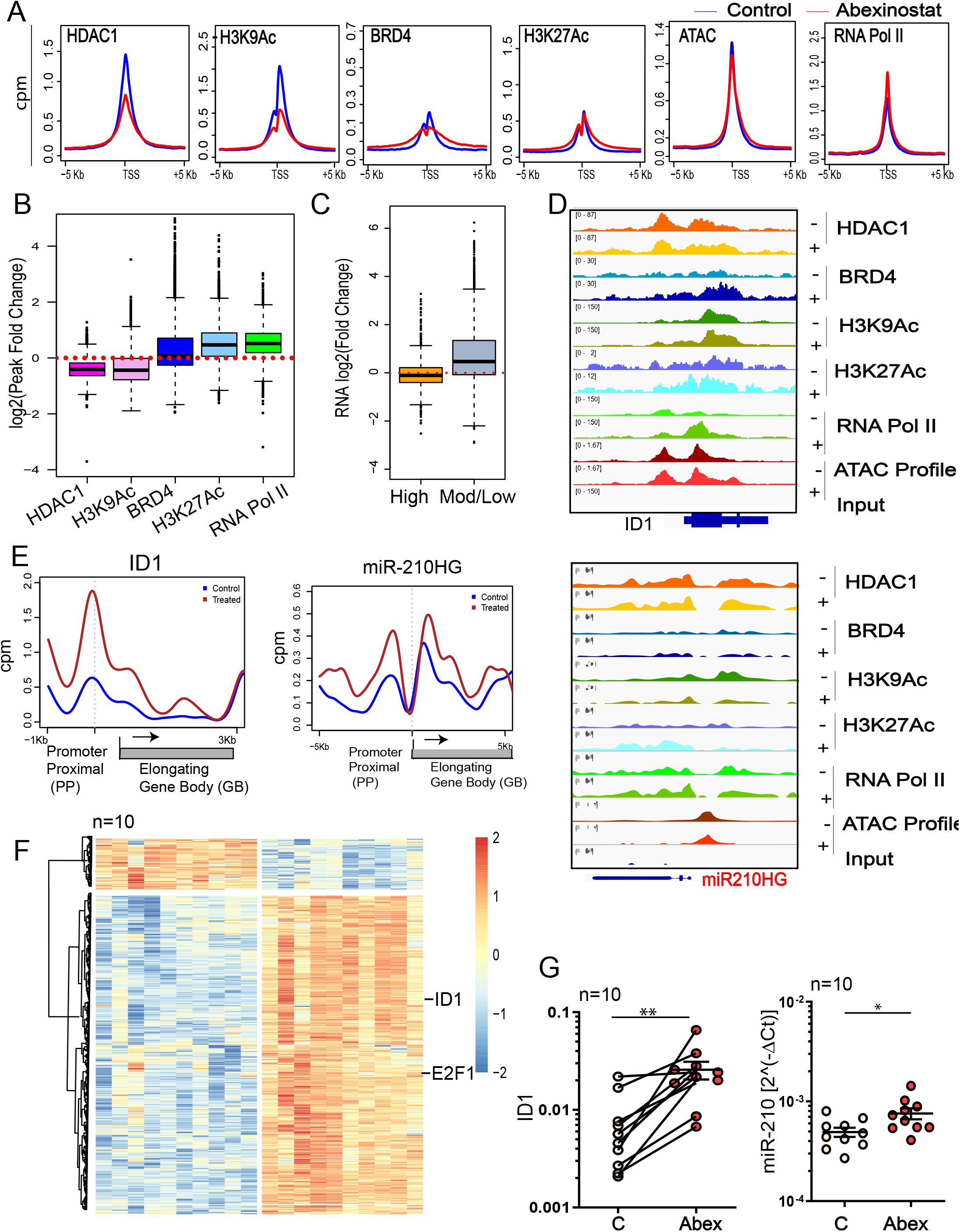
**(A)** Changes in the HDAC1, BRD4, H3K9Ac, H3K27Ac ATAC Seq and RNA Pol II ChIP-Seq meta-gene profiles before (blue) and after exposure (red) to the HDAC inhibitor, abexinostat for 6h at all 7958 peaks representing the union of HDAC1 co requited with H3K9Ac and Pol II without BRD4 at SEs (Cluster II). **(B)** Summary of changes in peak intensity (log2) for HDAC1, H3K9Ac, BRD4, H3K27Ac and Pol II after HDAC inhibition for 6h at all genes that had HDAC1 co requited with H3K9Ac and Pol II without BRD4 at SEs (Cluster II). **(C)** Based on average transcript expression at baseline, genes that bound HDAC1 at loci that also recruited H3K9Ac and showed open ATAC signatures and RNA Pol II were stratified into high (40%) or moderate/low (60%) expressers. mRNA Seq was conducted to assess the change in gene expression after exposure to abexinostat for 24 h and showed that the bulk of transcripts with low/moderate expression became upregulated. **(D)** Genome browser tracks showing the change in HDAC1 (orange), H3K9Ac (green), BRD4 (teal),H3K27Ac (dark Blue), RNA Pol II (green), ATAC Seq peaks (burgundy) in a representative CLL sample out of three at baseline and after exposure to abexinostat for 6 h, HDAC1 (yellow), H3K9Ac (navy), BRD4 (olive),H3K27Ac (sky Blue), RNA Pol II (dark green), ATAC Seq peaks (red) at each of the genes that bound HDAC1 at BRD4 dense SEs (Pax5, IKZF3, BLK,miR-155, Bcl2, BTK, HIST1H3C) and TEs (Syk and Ror1 and CD27. **(E)** Effect of HDAC inhibition on RNA Pol II initiation at promoters. Meta gene plots showing changes in reads per million (rpm) for RNA Pol II occupancy from −5 to −1KBkb of the proximal promoter (PP) centered around the transcriptional start site (TSS) to +3 to +6 KB of the gene body (GB) at each of the genes shown in (E) in CLL representative samples before (blue) and after exposure to abexinostat for 24 h (red). **(F)** RNA Seq of 10 CLL samples treated with the the HDAC inhibitor abexinostat (0.4 μM, 24h). Change in mRNA transcripts was assessed by RNA Seq compared with control samples; top 200 genes that showed a Log 2-fold change with p<0.05 of a total of 1000 protein coding genes that had HDAC bound across SEs and TEs **(G)** Validation of RNA Seq results using real time-PCR expression assays quantitating changes in expression levels of selected genes from Cluster II (ID1 and miR-210) from 10 CLL samples exposed to abexinostat for 24 h

### HDAC inhibition induces microRNA that have an inverse relation with driver genes in CLL

HDAC inhibition at loci where HDAC1 was bound either alone or with lowlevels of H3K9Ac, and without BRD4, H3K27Ac, open chromatin signatures or RNA Pol2 (Cluster III) elicited minimal loss of HDAC1 from target genes. in the genes in Cluster III responded to HDAC inhibition with strong hyperacetylation of both H3K9Ac and H3K27Ac (**Figure 7A**). Correspondingly, there was also a small but significant increase in BRD4 at target promoters presumably due to the increase in hyperacetylation as well as an increase in and RNA Pol2 engagement (**Figure 7A**); these changes are summarized in **Figure 7B**. At baseline over 80% of genes from Cluster III were expressed at moderate or low levels; HDAC inhibition induced over 80 % of these genes (**Figure 7C**); Changes in HDAC1, BRD4, H3K9Ac, H3K27Ac, open ATAC signatures and RNA Pol2 are shown for two representative microRNA, the miR-182 cluster and miR-9-3 (**Figure 7D**). Correspondingly, there was a significant increase in the RNA Pol2 engagement largely at the promoter and TSS indicating initiation of new transcription (**Figure 7E**).

**Figure 7.**
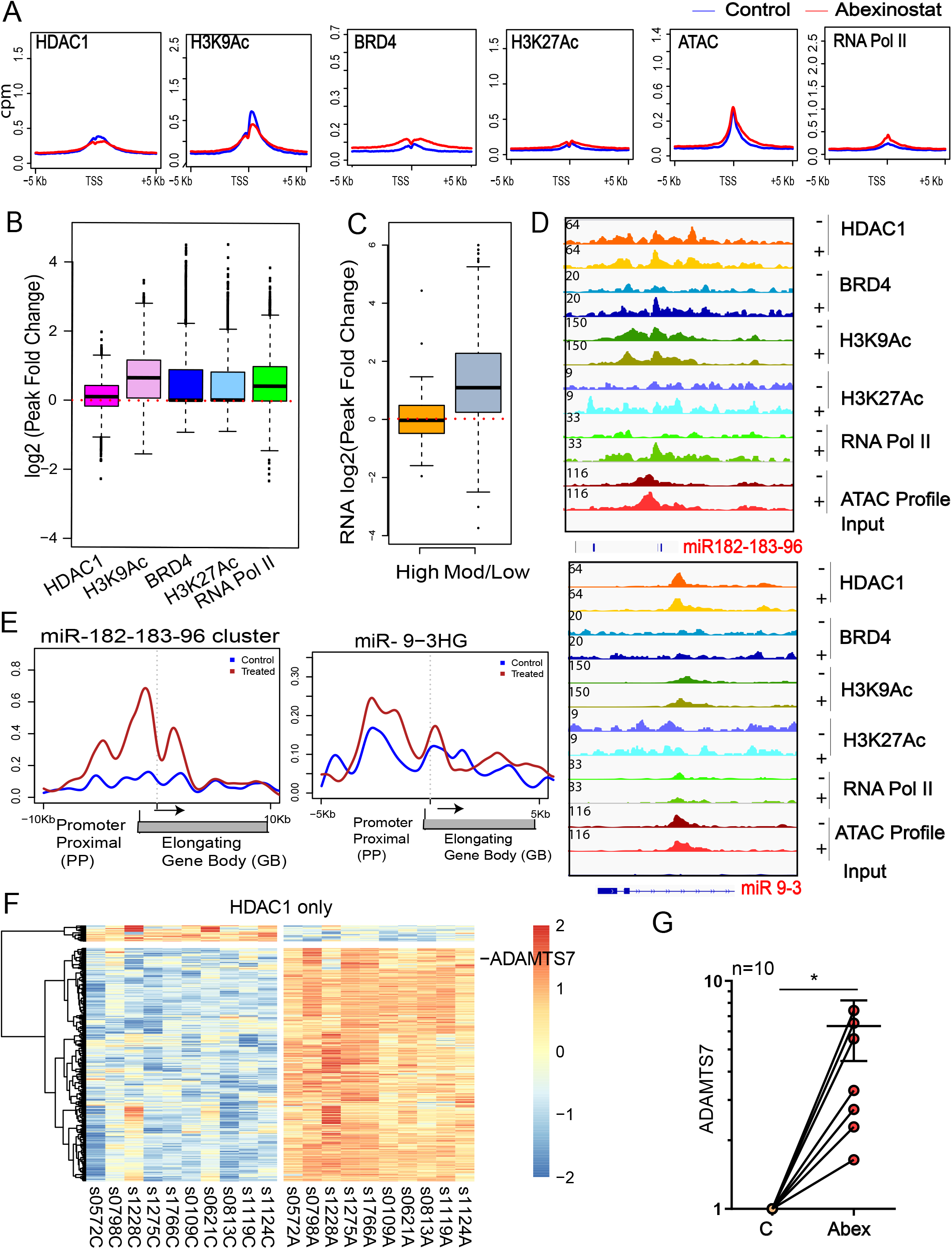
**(A)** Changes in the HDAC1, BRD4, H3K9Ac, H3K27Ac ATAC Seq and RNA Pol II ChIP-Seq meta-gene profiles before (blue) and after exposure (red) to the HDAC inhibitor, abexinostat for 6h at all 2271 peaks representing genes that recruited HDAC1 only without BRD4, SEs, H3K9Ac or RNA Pol II (Cluster III). **(B)** Summary of changes in peak intensity (log2) for HDAC1, H3K9Ac, BRD4, H3K27Ac and Pol II after HDAC inhibition for 6h at all genes that recruited HDAC1 only without BRD4, SEs, H3K9Ac or RNA Pol II (Cluster III). **(C)** Based on average transcript expression at baseline, genes that bound HDAC1 only without BRD4, SEs, H3K9Ac or RNA Pol II (Cluster III) were stratified into high (20%) or moderate/low (80%) expressers. mRNA Seq was conducted to assess the change in gene expression after exposure to abexinostat for 24 h and showed that 80% the bulk of transcripts with low/moderate expression became upregulated. **(D** Genome browser tracks showing the change in HDAC1 (orange), H3K9Ac (green), BRD4 (teal),H3K27Ac (dark Blue), RNA Pol II (green), ATAC Seq peaks (burgundy) in a representative CLL sample out of three at baseline and after exposure to abexinostat for 6 h, HDAC1 (yellow), H3K9Ac (navy), BRD4 (olive),H3K27Ac (sky Blue), RNA Pol II (dark green), ATAC Seq peaks (red) at representative genes (miR-182-183-96 cluster, miR-9-3) that recruited HDAC1 only without BRD4, SEs, H3K9Ac or RNA Pol II (Cluster III). **(E)** Effect of HDAC inhibition on RNA Pol II initiation at promoters. Meta gene plots showing changes in reads per million (rpm) for RNA Pol II occupancy from −5 to −1KBkb of the proximal promoter (PP) centered around the transcriptional start site (TSS) to +3 to +6 KB of the gene body (GB) at each of the genes shown in (E) in CLL representative samples before (blue) and after exposure to abexinostat for 24 h (red). **(F)** RNA Seq of 10 CLL samples treated with the the HDAC inhibitor abexinostat (0.4 μM, 24h). Change in mRNA transcripts was assessed by RNA Seq compared with control samples; top 200 genes that showed a Log 2-fold change with p<0.05 of a total of 1000 protein coding genes that had HDAC bound across SEs and TEs **(G)** Validation of ADAMTS7 expression from 10 CLL samples exposed to abexinostat for 24 h.

Next, we looked at changes in the transcriptome of both mRNA and small RNA. None of the protein coding genes induced in this cluster had significant roles in CLL biology (**Figure 7F, Table S8**); we validated and show the induction of ADAMTS7 as a representative protein coding gene (**Figure 7G**). When we evaluated the effect of HDAC inhibition on microRNAs, fold change rank ordering identified twelve microRNA that were differentially expressed in all 10 CLL samples with high significance (**Figure 8A**, **Tables S9,S10**); of which five miRNA belonged to Cluster III (miR-182 cluster, miR-9, miR-1303), four to Cluster II (miR210,miR-95,miR-92b,miR-320d) and one to Cluster I (miR-1248). Of these, the miR-182 cluster consisting of miR-182, miR183 and miR-96 were induced maximally (60-fold) whereas miR9-3 and all others were induced 4-fold or less. Induction of small RNA in primary tumor samples was validated in the TCL1 driven CLL mouse model which exposure to abexinostat led to the induction of miR-182, miR-183 and mR-96 (**Figure 8B, Table S11**). Small RNA expression assays confirmed the induction of these miRs in 10 CLL samples (**Figure 8C**). Since microRNA function by binding to and targeting mRNA for degradation or by preventing its translation, we co-sequenced microRNA and mRNA in the same CLL sample to identify potential miRNA-target gene interactions^29^. We focused on the miR-182 cluster as these were the microRNA with the highest induction. Bioinformatic analysis across twelve miRNA target prediction programs identified that all three microRNA in this cluster had binding sites on Pax5, IKZF3, BTK and Syk (**Table S12**). Anti-correlation analysis of miRNA and mRNA profiles in 10 CLL samples exposed to abexinostat show that induction of each of these miRNA was inversely correlated with declines in IKZF3 and BTK with high significance (rho >-0.55, P<0.05) whereas miR-182 and miR-183 were anti-correlated with Bcl2 and Syk (rho>-0.45, P<0.05) (**Figure 8D**). As a control we also evaluated the relation of these miRs with Pax5, Ror1, BLK and found that they were not significant (rho – p>0.05) (**Supp. Figure 5**) indicating that miR182-183 and 96 likely formed miRNA – mRNA target pairs with IKZF3,BTK, Bcl2 and Syk to decrease their expression.

**Figure 8.**
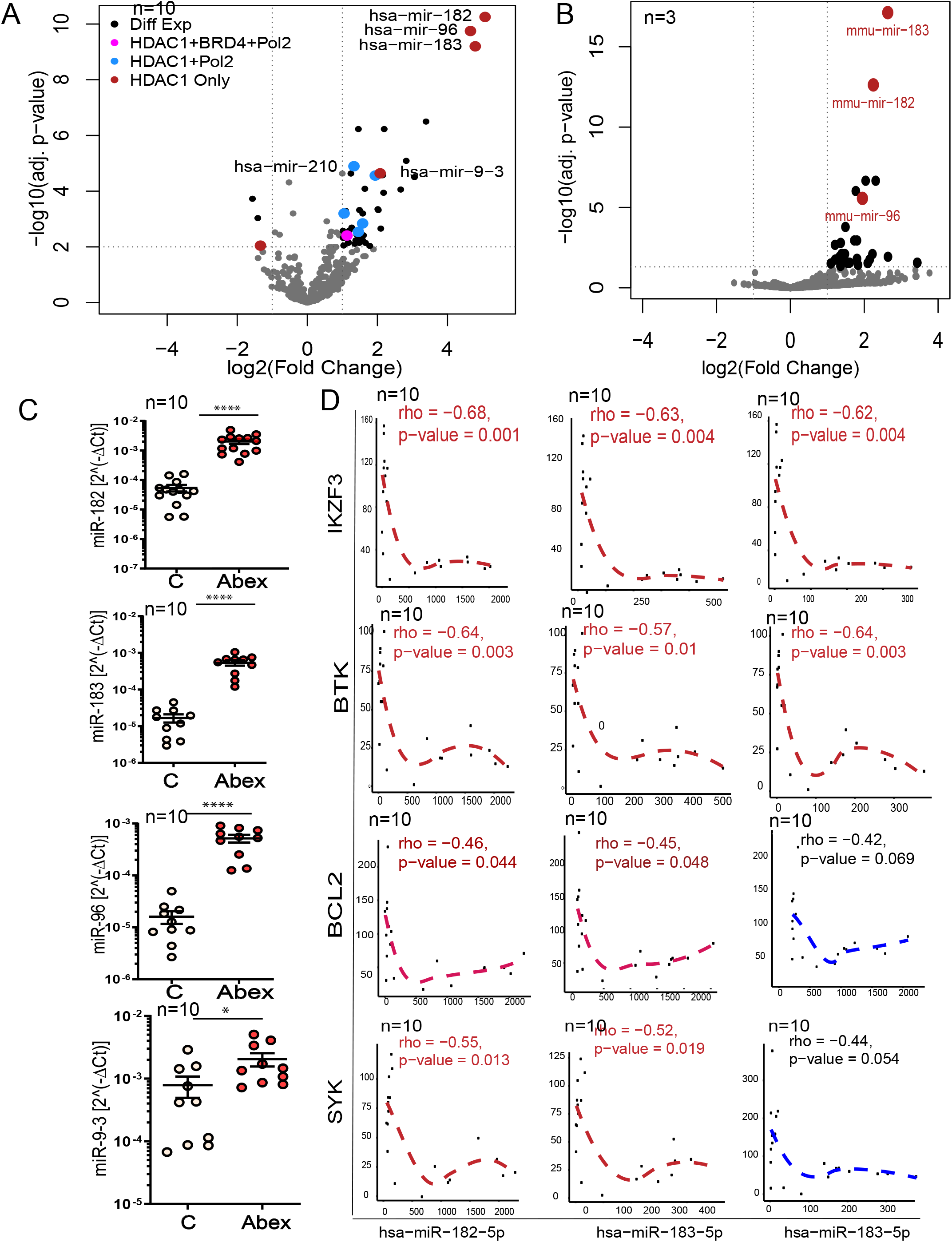
**(A)** Volcano plots showing the differential expression of small RNA genes assayed by small RNA Seq in 10 CLL samples before and after exposure to abexinostat for 24h. Of a total of ten microRNA genes that were significantly upregulated (adjust *P*-value<0.05 and log2 ratio>1),1 (miR-1224 in pink) belonged to Cluster I, five (miR-210, miR-95, miR92b, miR-320d, miR1296 in blue) to Cluster II and four (miR182-183-96 cluster, miR9-3 in red) to Cluster III. One gene (miR-1303) from Cluster III was downregulated (adjust *P*-value<0.05 and log2 ratio<–1). **(B)** Volcano plots showing the differential expression of small RNA genes assayed by small RNA Seq in 3 samples obtained from Tcl-1 mice with CLL randomized to treatment with vehicle or abexinostat for two weeks. The top three micrpRNA that became significantly induced (adjust *P*-value<0.05 and log2 ratio>1) in all three samples are marked in red. **(C)** mRNA expression levels of genes from Cluster III (miR-182, miR-183, miR-96 and miR9-3) from 10 CLL samples exposed to abexinostat for 24h. **(D)** The Pearson correlation between the top microRNA induced (miR-182, miR-183 and miR-96) in both primary CLL samples and TCl-1 bearing mice were evaluated for negative correlation against their predicted target driver genes relevant to CLL (IKZF3, BTK, BCL2, and Syk). We show the Pearson scatter plots showing a negative correlation between miR-182, miR-183 and miR-96 and IKZF3 (p value = 0.001 for miR-182, 0.003 for miR183 and 0.013 for miR-96), BTK (p value = 0.004 for miR-182, 0.01 for miR183 and 0.01 for miR-96),Bcl2 and Syk (p value = 0.004 for miR-182, 0.003 for miR183 and 0.054 for miR-96). Rho values and slopes are shown.

## DISCUSSION

The histone deacetylase HDAC1 has been extensively characterized as a transcriptional repressor in CLL and across cancers. However, recent reports also described it as necessary for core transcription factor expression and as a transcriptional activator at super-enhancers. Our present study provides an extensive characterization of the functional epigenome in CLL, extends previous studies, and is the first to comprehensively characterize the context under which HDAC1 governs the regulatory landscape to facilitate and repress individual target genes within the same tumor to dysregulate the transcriptome and favor tumor survival.

Previous attempts to establish the regulatory epigenome in CLL focused on the differences between normal B cells and CLL. This work measured gain or loss of enhancers based on their H3K27Ac signal and related them to binding motifs of transcriptional factors in CLL and B cells but did not experimentally verify binding of identified transcription factors at their predicted targets or link chromatin states to transcriptional profiles^30^. Our group and others previously profiled the chromatin reader BRD4 and found that BRD4 was associated with specific SEs and controlled the expression of key oncogenic proteins in CLL^17^ as well as drove the transcription of Pax5, a key transcriptional factor required for CLL survival^22^. Targeting BRD4 led to a loss of SEs and oncogenic transcription^17,22^. These findings identified the drivers of oncogenic networks but do not address the vast fraction of the transcriptome that are not bound by BRD4 or H3K27 but play an important role in cancer.

The role of HDACs in transcription has not been clearly elucidated yet. Some reports identify that HDAC1 is required for the active transcription of key transcription factors in rhabdomysarcoma and showed that HDAC1/2/3 were associated with SEs in these cell lines and facilitated transcription^18,24^. HDAC inhibition in this setting caused hyperacetylation of H3K27 initially followed by collapse of SE and H3K27Ac within 6 h and loss of SE driven transcripts. Yet others suggest that HDAC is required for transcriptional elongation not initiation^25,31^. HDAC1 was also shown to regulate BRD4 availability; HDAC inhibition in this cell line model was proposed to trap BRD4 at hyperacetylated histones sequestering it and preventing its redistribution to newly induced genes preventing induction^31^.

Our study shows that HDAC1 has a complex but central role in the regulatory chromatin landscape of CLL and drives survival by facilitating the transcriptional dysregulation characteristic of this disease. When we conducted genome wide studies of functionally related, non-overlapping chromatin modifications and integrated the recruitment of HDAC1, BRD4, acetylated H3H9 and H3K27, and RNA Pol2 engagement with chromatin accessibility and expression analysis we found that HDAC1 is co-recruited with BRD4 at SEs marked by hyperacetylated H3K27. These loci exhibited highly acetylated H3K9Ac, open chromatin signatures and RNA Pol2 engagement and the bulk of genes in this cluster (Cluster I) were robustly expressed. Although we saw that BRD4 was recruited to a slightly larger subset of chromatin loci without accompanying HDAC1 recruitment, it became clear most of the genes that were important in the pathology of CLL were bound by BRD4 and HDAC1 and that HDAC1 functioned as transcriptional activator at BRD4 marked SEs in CLL. Largely, genes critical to the pathology of CLL such as the transcription factor Pax5, IKZF3, members of the BCR survival pathway, several immune regulators such as CXCR4, IL16, IL4R and the anti-apoptotic proteins Bcl2, belonged to Cluster I. Mechanistic evidence supporting a critical role for HDAC1 in the activation of these genes comes from results where HDAC inhibition causes a steep decline in the expression of these driver genes in CLL. While HDAC inhibition was as effective as targeting BRD4 in attenuating driver gene expression, we found critical differences in their mechanism of action. BRD4 inhibitors caused a clear decrease in SEs at driver genes and decreased expression of target genes perhaps by reducing eRNA production, but the link between BRD4 inhibition and decreases in RNA Pol2 at target promoters was unclear^17,22^. In contrast to the response elicited by both BRDi and HDAC inhibition in rhabdomyoma cell lines, HDAC inhibition in tumor CLL cells did not decrease H3K27AC levels. Instead, it selectively decreased BRD4 binding at promoters and disrupted RNA Pol2 binding at transcriptional initiation sites, effectively decreasing the expression of multiple CLL driver genes. In addition to driver protein coding genes, this cluster included the oncogenic microRNAs, miR-155 and miR-21. miR-155 affects BCR signaling^32^ and both miR-155 and miR-21^33^ were shown to be associated with adverse outcomes in CLL by us and others. Significantly, HDAC1 was not found in a complex with BRD4, indicating that they each exerted independent effects on the regulatory chromatin and that the consequence of HDAC inhibition on transcription was not due to effects on a co-complexed BRD4. These results clearly establish that the importance of HDAC1as an independent regulator that is critical in driving the expression of oncogenic driver genes in CLL.

However, HDAC1 has roles beyond transcriptional activation in CLL. Our study also delineates the configuration of the regulatory chromatin landscape that enables HDAC1 to function as a transcriptional repressor. When HDAC1 is recruited without BRD4 or SEs at chromatin loci it acts as a transcriptional repressor; a set of these loci (Cluster II) display high levels of H3K9Ac and RNA Pol2 engagement. Despite this, the bulk of genes in this cluster were expressed at moderate or low levels (60%) and included protein coding genes as well as several microRNA such as miR-210, miR-95 and miR320. HDAC1 was also recruited to chromatin loci without BRD4 and SEs; these loci also lacked acetylation at H3K9Ac, open ATAC signatures and RNA Pol2 engagement (Cluster III) and over 80% of targets in this cluster were silenced. Cluster III did not include protein coding genes relevant to CLL it did include many key microRNA such as the miR-182-183 and 96 cluster as well as miR 9-1-2 and 3. Largely, HDAC inhibition served to induce genes belonging to Clusters II and III indicating that in the absence of association with SEs, HDAC1 functions as a transcriptional repressor.

MicroRNA have long been known to be dysregulated in CLL and closely involved in the pathology and progression of this disease^5,6^. Of the 10 microRNA that showed statistically significant induction in all 10 CLL samples after HDAC inhibition, four of these belong to Cluster II and five belonged to Cluster III. The microRNA such as miR-210 has been previously validated by our group^2^. HDAC inhibition induced miR-210 to downregulate BTK, a protein central to CLL survival. The importance of miRs in regulating proteins of critical importance in CLL such as BTK is supported by results in this paper that showed that BTK transcripts were unaffected by HDAC inhibition and did not lose RNA Poll II recruitment. Rather, the steep declines in BTK protein likely reflected miRNA targeting by mir-210 and the miR-182 cluster identified in this study. We observed that the miR-182-183 and 95 cluster was induced between 35-60 fold indicating that they were exquisitely responsive to HDAC inhibition. In this study we used co-sequencing of microRNA and mRNA within the same samples to conduct inverse-correlation analysis of miRNA and mRNA expression data to identify predicted functional miRNA targets. This method allows us to evaluate relationships in intact tumor cells, and is likely to be more physiologically representative compared to methods that rely on overexpression of miR followed by immunoblotting as previously shown by us^2,8^ or miR-mRNA reporter based^2,8^ and sequencing based analysis^34^. miR targeting of mRNA is context dependent and depends on the expression of relevant targets and absence of competing RNA making the use of cell lines which do not recapitulate the expression patterns of primary CLL cells for such experiments less relevant. Our study showed that the miR-182-183-96 cluster expression after HDAC inhibition was significantly anti-correlated with expression of Pax5, IKZF3 and BTK each of which was a critical regulator of survival in CLL.

When we validated key findings from our primary CLL samples in vivo using an Eμ-TCL1 mouse model of CLL, we found that abexinostat treatment led to a strong declines in the levels of a set of genes shared in human CLL such as decreases in BTK, BLK, PAX%, IKZF3, IKZF1, CD37, CD79b and increase in HIST genes as well as miR-182, miR-183 and miR-96.

In conclusion, our study indicates that HDAC1 functions both as an activator by selectively associating with BRD4 at SEs and facilitating the transcription of driver genes in CLL or as the repressor by associating independently at target loci to silence the expression of key microRNA that in turn allows the deregulated abundance of hallmark survival genes in CLL. These complimentary actions establish a transcriptional dependency by facilitating the abundant aberrant expression of CLL specific transcriptional factors, survival and signaling proteins that are critical to the biology and survival of CLL cells.

